# Temperature-based MHC class-I multimer peptide exchange for human HLA-A, B and C

**DOI:** 10.1101/2024.12.23.630039

**Authors:** Cilia R. Pothast, Ian Derksen, Anneloes van der Plas – van Duijn, Angela el Hebieshy, Wesley Huisman, Kees L.M.C. Franken, Jacques Neefjes, Jolien J. Luimstra, Marieke Griffioen, Michel Kester, Maarten H. Vermeer, Marcus Östholm, Sine R. Hadrup, Mirjam H.M. Heemskerk, Ferenc A. Scheeren

## Abstract

T cell recognition of specific antigens presented by major histocompatibility complexes class-I (MHC-I) can play an important role during immune responses against pathogens and cancer cells. Detection of T cell immunity is based on assessing the presence of antigen-specific cytotoxic CD8+ T cells using MHC class-I (MHC-I) multimer technology. Previously we have designed conditional peptides for HLA-A*02:01, H-2K^b^ and HLA-E that form stable peptide-MHC-I-complexes at low temperatures and dissociate when exposed to a defined elevated temperature. The resulting conditional MHC-I complex can easily and without additional handling be exchanged with a peptide of interest, allowing to exchange peptides in a ready-to-use multimer and a high-throughput manner. Here we present data that this peptide-exchange technology is a general applicable, ready-to-use and fast approach to load many different peptides in MHC-I multimers for alleles of the HLA-A, HLA-B and HLA-C loci. We describe the development of conditional peptides for HLA-A*03:01, HLA-A*11:01, HLA-B*07:02 and HLA-C*07:02 that only form stable peptide-MHC-I complexes at low temperatures, allowing peptide exchange at higher defined temperature. We document the ease and flexibility of this technology by monitoring CD8+ T cell responses to virus-specific peptide-MHC complexes in patients.

**Highlights:**

- T cell immunity relies on antigen-specific CD8+ T cells recognizing peptide MHC-I complexes.
- Establishing temperature-based peptide exchange across multiple HLA alleles, resulting in a robust, easy, and fast system to generate peptide MHC-I complexes.
- Temperature-based MHC class-I multimer demonstrate applicability across major MHC-I gene families for monitoring CD8+ T cell responses.
- Easy high-throughput peptide exchange potential, enhancing clinical utility of MHC multimer technology.

**Graphical abstract:** 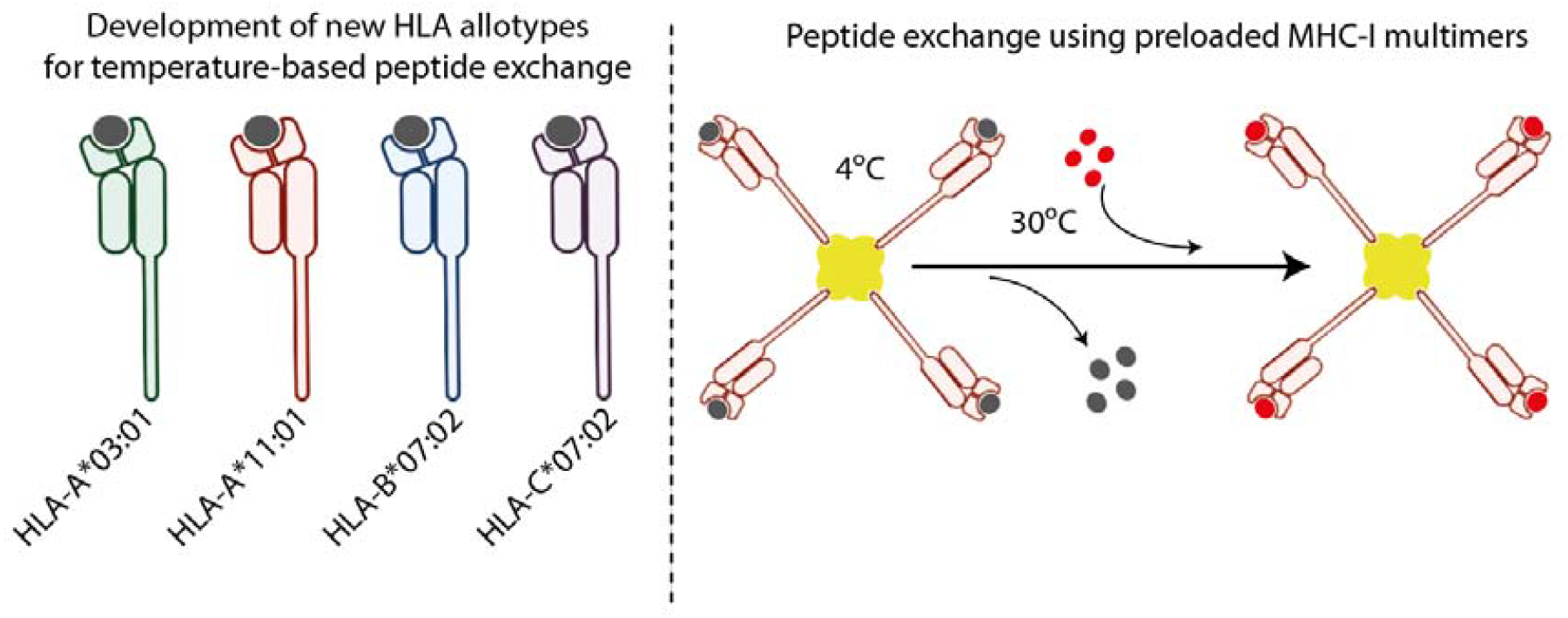

## Introduction

An essential component of cellular immunity is mediated through the T Cell Receptor (TCR) of CD8+ T cell that can bind MHC-I complexes. The MHC-I complex consist of three components: a polymorphic heavy chain, the β2-microglobulin (β2M) protein, and an 8–10-amino-acid peptide ligand[1]. Peptide selection occurs through docking of the peptide C-terminus in MHC-I and occupation of the pockets in MHC-I followed by a conformational change that preferably traps the peptides with highest MHC-binding affinity. As a result, peptides with low MHC-binding affinity are replaced by peptides with a high MHC-binding affinity, a phenomenon used in this work [2]. The peptide ligand is an essential part of the MHC-I complex; without it, the complex cannot remain stable and dissociate[3]. Pathogens, such as viruses, can infect human cells and a primary infection often leads to disease and immunity. Recurrent infections typically do not cause severe disease and are cleared, in part by T cells recognizing viral peptides presented by MHC-I, as recently demonstrated for SARS-CoV-2[4, 5]. The emergence of viral variants can potentially give rise to a partial change in immunogenic viral peptides thereby affecting disease progression. Moreover, immune-compromised patients who lack proper B cell immunity are fully dependent on T cell-mediated immunity. Therefore, monitoring antigen-specific T cells is crucial for tracking the impact of recurrent infections and determining whether therapeutic intervention is required.

The detection and visualization of antigen-specific T cells was first achieved by using peptide-loaded MHC-I multimers with a fluorochrome[6]. To generate such labelled multimers, soluble MHC-I monomers complexed with a peptide are multimerized through biotinylating and fluorochrome-conjugated streptavidin. This process is time-consuming and involves multiple steps for each individual peptide-MHC-I complex. Several technologies have since been developed to allow the parallel production of multiple MHC-I multimers, facilitating high-throughput analysis. For instance, dipeptides have been used as catalysts for peptide exchange[7-10], UV-sensitive conditional MHC ligands have been a widely adopted approach[11, 12], in which a photocleavable peptide is cleaved upon UV exposure. Additional peptide exchange technologies include: chaperone-mediated peptide exchange[13, 14] enzymatic cleavage of low affinity peptide that are covalently linked [15] and small alcohol peptide exchange [16]. This results in an unloaded MHC-I monomer, which can subsequently be loaded with an high-affinity peptide and multimerized[11, 12]. However, UV exposure can cause photodamage to the peptide MHC-I complex, that leads to sample evaporation, and is incompatible with fluorescently labeled multimers, requiring multimerization to be performed after the peptide exchange[17]. Additionally, more recent methods involve empty MHC-I stabilized by the introduction of a disulfide bond, which can be loaded with the desired a peptide of choice. It is currently unknown how these mutations and structural alterations affect the peptide binding repertoire of these disulfide-stabilized MHC-I molecules[8, 9].

One major limitation of current MHC-I multimer-based technologies is the complex, multistep protocols to obtain ready-to-use peptide MHC-I complexes, without inducing mutations in the HLA class I heavy chain or B2M. This limits the broad clinical usage of MHC-I multimer technology. To address this, we previously developed a temperature-based peptide exchangeable MHC-I multimer technology for human HLA-A*02:01, HLA-E and murine H2Kb, which simplifies and accelerates peptide exchange on MHC-I multimers[18-20]. This technology allows the generation of multimers within hours with ease, simply by incubating selected peptides at a defined temperature with a single standardized batch of fluorescently labeled MHC-I multimers [21]. Depending on the platform, peptide exchange may be performed either before or after multimerization and labeling. In this study, we aimed to determine whether this technology could be developed into a robust and broadly applicable system for analyzing CD8+ T cell immunity. To this end, we developed temperature-based peptide-exchange MHC-I multimers for human MHC gene product HLA-A*03:01, HLA-A*11:01, HLA-B*07:02, and HLA-C*07:02, representing all major MHC-I gene families.

## Results

### Design of temperature-exchangeable template peptide

We set out to extend our toolbox of HLA alleles that allow temperature-based template peptide exchange using multimerized, ready-to-use MHC-I complexes. Our goal was to design peptides that form stable complexes with MHC-I at low temperature and are unstable at higher temperatures, allowing replacement of the template peptide by a desired high-affinity peptide. The peptide off-rate is the primary determinant for the MHC-I stability[22]. For HLA–A*03:01, we designed peptides based on a Hepatitis C virus epitope (HCV43-51; RLGVRATRK) (dissociation constant [Kd] 11.7 nM)[23] and modified the residues to decrease the dissociation constant[24]. HLA–A*03:01 complexes were produced with wild-type or modified peptide, and thermal stability was assessed at 50°C using analytical size exclusion high-performance liquid chromatography (HPLC). With the modified peptide, RGAVRATRR (Kd 7098.44 nM), the complex was unstable at 50°C, resulting in denaturation, as shown by the absence of a MHC-I peak in a size-exclusion HPLC (Figure 1 & Supplemental Figure 1). In contrast, the presence of the high-affinity epitope (RLGVRATRK, 11.7 nM) produced a distinct peak, demonstrating that the HLA-A*03:01 complex could be rescued from unfolding. A similar approach was applied to HLA-A*11:01, HLA-B*07:02 and HLA-C*07:02. For each allele, we designed peptides for each respective allele and modified the residues to decrease the affinity (Table 1)[23, 25, 26]. HLA complexes with the modified peptides were produced, and their thermal stability was evaluated at 50°C using analytical size-exclusion HPLC. For each allele, we identified a thermal template peptide that was unstable at 50°C, which could be rescued by the addition of a high-affinity peptide (Figure 1 & Supplemental Figure 1). We proceeded with peptides that exhibited thermo-instability at 50°C that showed a rescued HLA complex in the presence of the high-affinity peptide. The peptides that formed stable complexes at 4⍰°C but unstable complexes at 20–32⍰°C and 50⍰°C displayed affinities ranging from 4 to 11⍰μM. Such low affinities are uncommon for endogenous peptides and are unlikely to have physiological relevance, given that the complexes are highly unstable at 37⍰°C.

**Table 1:** Peptide used for template development

| MHC-I allele | Source | Epitope | Sequence | K<sub>d</sub> (nM) |  |
| --- | --- | --- | --- | --- | --- |
| A*03:01 | Hepatitis C virus | HCV<sub>43-51</sub> | RLGVRATRK | 11.7 | [24] |
|  |  |  | R**GA**VRATR**R** | 7098.44 | [23] |
|  |  |  | R**AG**VRAT**KR** | 8206.05 | [23] |
|  |  |  | R**SG**VRATR**H** | 8293.07 | [23] |
|  |  |  | R**GA**VRAT**KR** | 8654.53 | [23] |
|  |  |  | R**AG**VRATR**H** | 13381.76 | [23] |
| A*11:01 | Homo sapiens | PLA-X<sub>26-34</sub> | KVVNPLFEK | 8.1 | [25] |
|  | Homo sapiens | GAD-65<sub>255-263</sub> | RFKMFPEVK | 3898.11 | [23] |
|  |  |  | R**R**KMFPEVK | 21628.45 | [23] |
|  |  |  | RFKMFPEV**R** | 11988.30 | [23] |
|  | Dengue virus,<br>Genome polyprotein | 2007-2015 | KTFVELMRR | 30.08 | [23] |
|  |  |  | K**AF**VDLMR**H** | 3871.92 | [23] |
|  |  |  | KTFVDL**GRH** | 3277.65 | [23] |
|  |  |  | KTFVDL**ARH** | 1659.70 | [23] |
|  |  |  | KTFVDLMR**H** | 414.23 | [23] |
| B*07:02 | Homo sapiens | AKR1C3<sub>91-99</sub> | RPELVRPAL | 4.38 | [23] |
|  |  |  | **LA**ELVRPAL | 732.89 | [23] |
|  |  |  | **VA**ELVRPAL | 826.71 | [23] |
|  |  |  | R**K**ELVRPAL | 4118.37 | [23] |
| C*07:02 | M. tuberculosis | EsxH<sub>4-12</sub> | IMYNYPAML | 67 | [26] |
|  |  |  | **I**AYNYPAM**A** | 5158.68 | [23] |
|  |  |  | **I**KYNYPAM**A** | 9139.38 | [23] |

**Figure 1:**
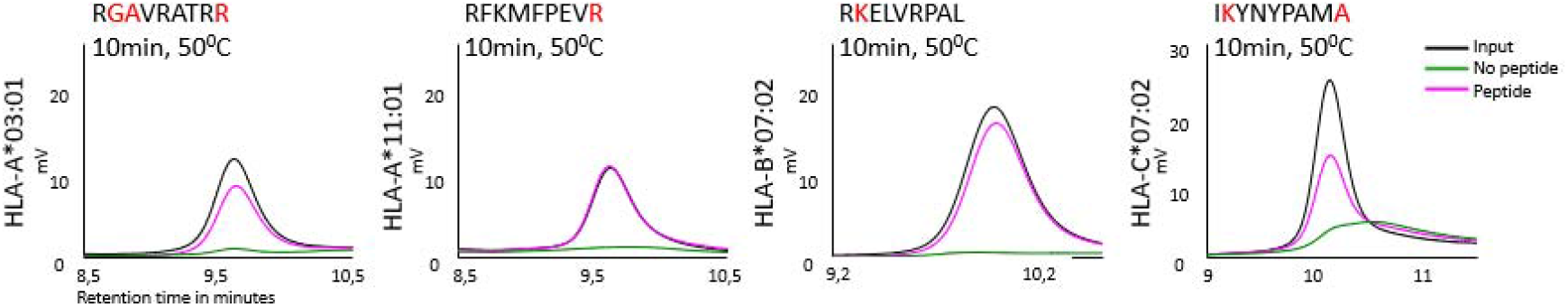
Thermal stability of MHC-I monomers with template peptide. Thermal stability of HLA-A03:01, HLA-A11:01, HLA-B07:02, HLA-C07:01, and HLA-C*07:02 complexed with their template peptides was assessed at 50⍰°C using analytical size-exclusion HPLC. Peptide–MHC I monomers were incubated with indicated peptides at 50°C for 10 minutes. Peptides used are listed in Table 1. One of at least three representative experiments is shown. The black line represents primary data without thermal exchange, the green line shows thermal exchange without peptide (resulting in complex instability), and the pink line represents thermal exchange in the presence of peptide (yielding a stable complex).

Next, we optimized the temperature and time for peptide exchange for each allele. For HLA– A*03:01, we tested three template peptides that were unstable at 50°C and could be rescued in the presence of a high-affinity peptide. Thermal stability was assessed by incubating the HLA–A*03:01 complex with a high-affinity peptide (RGAVRATRR) at various temperatures for 1 hour (Figure 2 & Supplemental Figure 2). We selected the template peptide RGAVRATKR, which had the lowest optimal exchange temperature (23°C) for 1 hour. This condition resulted in a nearly complete unfolding of the complex in the absence of a high-affinity peptide and full rescue of the HLA– A*03:01 complex in the presence of a high-affinity peptide (Figure 2). Similar results were obtained when the complexes were incubated at 30°C. A similar approach was used for HLA-A*11:01, HLA-B*07:02, and HLA-C*07:02, where the optimal peptide exchange conditions entailed an 1-hour incubation at 30°C (Figure 2 & Supplemental Figure 2). The exchange efficiency was then examined in detail for all 4 of the alleles included in this study. We found that replacing the template peptide with the high-affinity peptide and incubating for approximately 1–1.5 hours at the specified temperature resulted in nearly complete peptide exchange, as shown in Figure⍰2B. This pattern was consistent across the different HLA alleles tested and aligns with the high exchange efficiencies previously documented for HLA-A*02:01 and H2Kb[18]. For 3 alleles, the measured exchange efficiency exceeded 100%. This effect arises from the intrinsic instability of the input complex at temperatures above 4°C, which partially dissociates during the short preparation and measurement steps. In the condition when a high affinity peptide is present the peptide stabilizes the complex. Thus, the efficiency of thermal exchange may be underestimated due to technical issues. Collectively, these findings establish robust conditions for efficient, thermal peptide exchange across multiple HLA class I alleles and expand the applicability of thermal peptide exchange for multimer production.To verify the quality of the thermally exchanged peptide–MHC (pMHC) complexes, we incorporated an additional control based on LILRB1 binding. Stability of each pMHC complex was assessed using K562 cells retrovirally transduced to overexpress LILRB1 (Fig.⍰2C). Because LILRB1 recognizes structural elements of β2⍰microglobulin (β2M) and the α3 domain of the MHC heavy chain, it selectively interacts with properly assembled and stable pMHC complexes. We have previously demonstrated this for both thermally exchanged HLA⍰E tetramers and UV⍰mediated peptide exchange reagents [27, 28]. Consistent with this, four conventional PE⍰conjugated tetramers stained only K562-LILRB1^OE^ cells (Fig.⍰2D). We then applied the thermal peptide⍰exchange tetramers, representing four HLA alleles. When a high affinity peptide formed a stable complex with the PE⍰conjugated tetramer, robust staining of K562-LILRB1^OE^ cells was observed (Fig.⍰2E). In the absence of peptide, unstable association with the MHC heavy chain led to β2M dissociation from the fluorescent tetramer and no LILRB1-dependent staining was observed. Collectively, these optimizations and validation steps confirm that thermal peptide exchange is a robust and versatile method for generating stable pMHC complexes, enabling reliable downstream detection of antigen-specific T cells.

**Figure 2:**
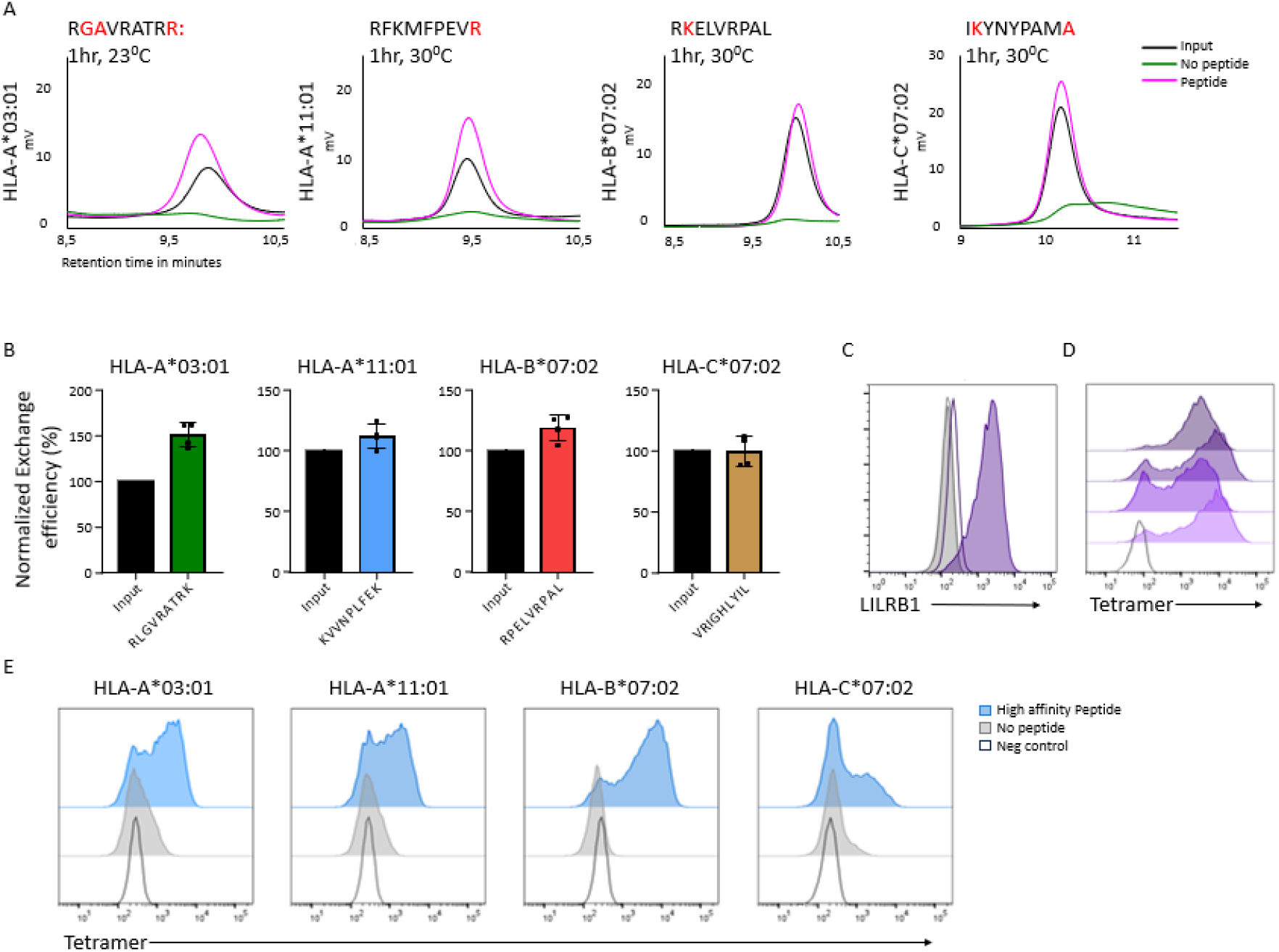
Optimal exchange conditions for template-peptide MHC-I monomers. A) Thermal stability of the template peptide-MHC-I complex to determine optimal exchange temperature with one high-affinity peptide for all alleles tested. Primary data of temperature-based peptide exchange analyzed by gel filtration chromatography at indicated temperatures. The black line represents primary data without thermal exchange, the green line shows thermal exchange without peptide (resulting in complex instability), and the pink line represents thermal exchange in the presence of peptide (yielding a stable complex). The thermal exchange temperatures corresponding to each histogram are indicated above the panels and are 23, 30, 30, and 30⍰°C, respectively. Peptides used are depicted in Table 1. One of at least three representative experiments is shown. B) The exchange efficiency was calculated from the area under the curve measured by HPLC and normalized to binding of the optimal peptide and timing as indicated in each histogram. Mean values ± SD from three independent experiments are shown. C) LILRB1 over-expression (OE) in K562. Grey: negative control, Purple line: K562-WT plus anti-LILRB1-Fitc, Purple filled: K562-LILRB1^OE^ plus anti-LILRB1-Fitc (N=3). D) 4 different conventional tetramer staining on K562-LILRB1^OE^. Open blackline: negative control. (N-6) E) thermal exchange using a high affinity peptide or no peptide, followed by staining of the K562-LILRB1^OE^ and flow cytometric analysis (N=4).

### Temperature-exchanged MHC-I multimers for detection of antigen-specific CD8+ T cells

Previously, we demonstrated that HLA-A*02:01 MHC-I multimers generated via temperature-based peptide exchange performed comparable to those produced using conventional methods[18, 21]. We also established that peptide exchange can be directly applied to multimers themselves, resulting in a more user-friendly protocol with reduced handling time[18, 21]. To evaluate the effectiveness of the newly developed temperature-exchanged MHC-I multimers for detection of antigen-specific T cell responses, we performed temperature exchange reactions using a panel of high-affinity peptides specific for HLA-A*03:01, HLA-A*11:01, HLA-B*07:02, and HLA-C*07:02. These temperature-exchanged MHC-I multimers efficiently labelled clonal T cell lines (Figure 3), in a concentration dependent manner (Supplemental Figure 3). This evaluation included two HLA-A*03:01-restricted T cell lines, one HLA-A*11:01-restricted T cell line, two HLA-B*07:02-restricted T cell lines, and one HLA-C*07:02-restricted T cell lines. A complete list of peptides used for temperature-based peptide exchange is provided in Supplemental Table 2. We then compared the staining of T cells by temperature-exchanged MHC-I multimers to those stained with classical folded MHC-I multimers. In the latter case, each MHC-I multimer has to be produced individually, while temperature exchange allows for a batch that can be taken from the freezer and warmed up in the presence of intended peptides before flow analyses. This procedure takes only 1 hr. For this comparison, we generated temperature-exchanged MHC-I multimers for each of the four newly developed alleles and compared their staining performance to that of the classical folded MHC-I multimers at isomass for the same peptide (Figure 4). Additionally, staining intensity was evaluated across multiple dilutions using isomass MHC-I multimer concentrations (Supplemental Figure 4). Overall, the temperature-exchanged MHC-I multimers showed distinct binding to clonal T cells in an antigen-specific manner and exhibited only a minor decrease in performance compared to the classical MHC-I multimers, particularly for the HLA-A11:01 and HLA-B07:02 complexes. Temperature-exchanged multimers for HLA-A*03:01 and HLA-C*07:02 showed similar or slightly lower staining intensities compared to their classical counterparts. Higher concentrations of both multimer types can be used to further enhance staining intensity without any technical complications.

**Figure 3:**
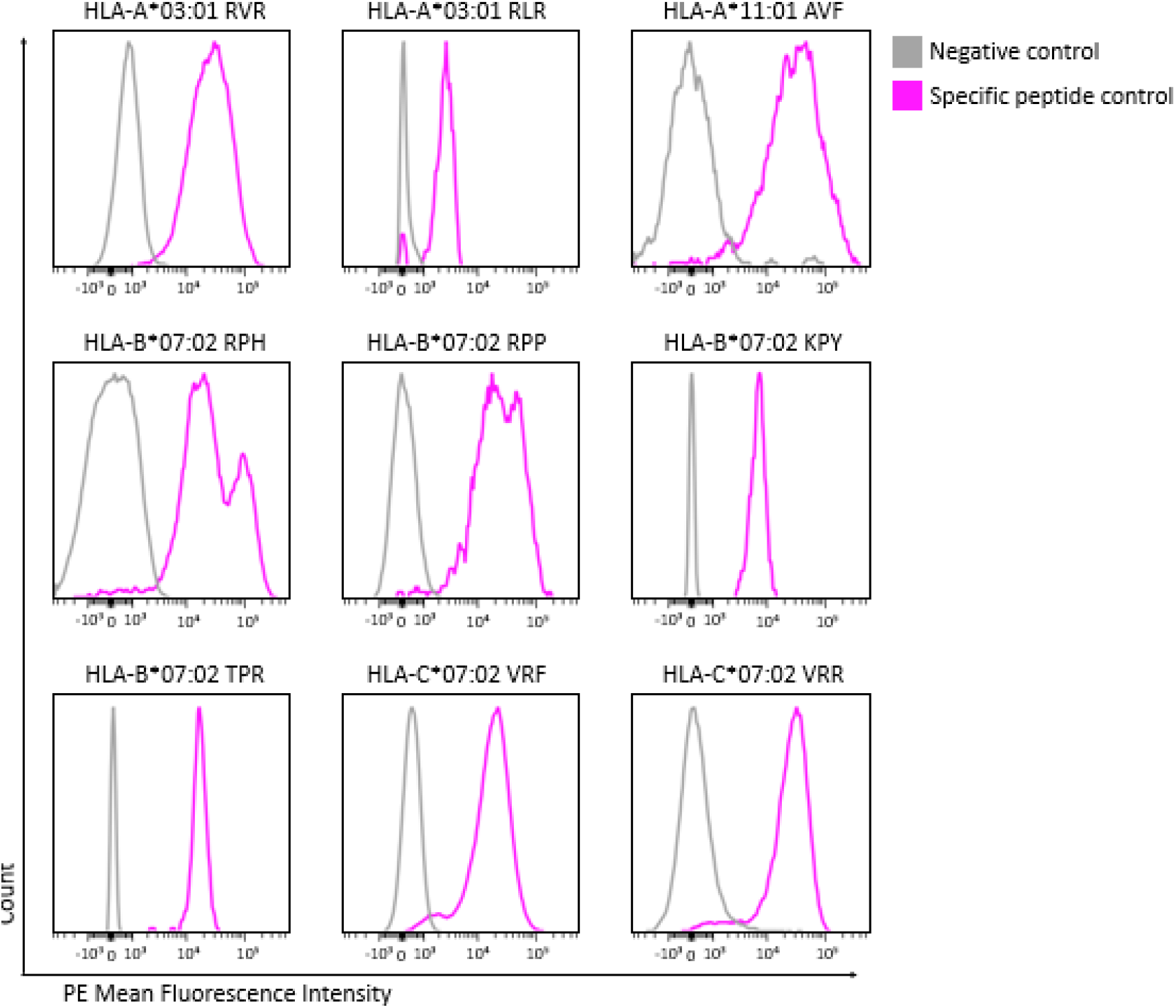
Temperature exchange tetramer binding to clonal T cell lines. Clonal CD8+ T cell lines stained with PE-conjugated temperature exchange multimers that either contained a relevant (pink) or irrelevant peptide (grey). The specificities of the T cell lines are depicted above each figure, as well as HLA-type and first three amino acids of the peptide. Y-axis shows the individually scaled counts and x-axis shows the mean fluorescent intensity of PE.

**Figure 4:**
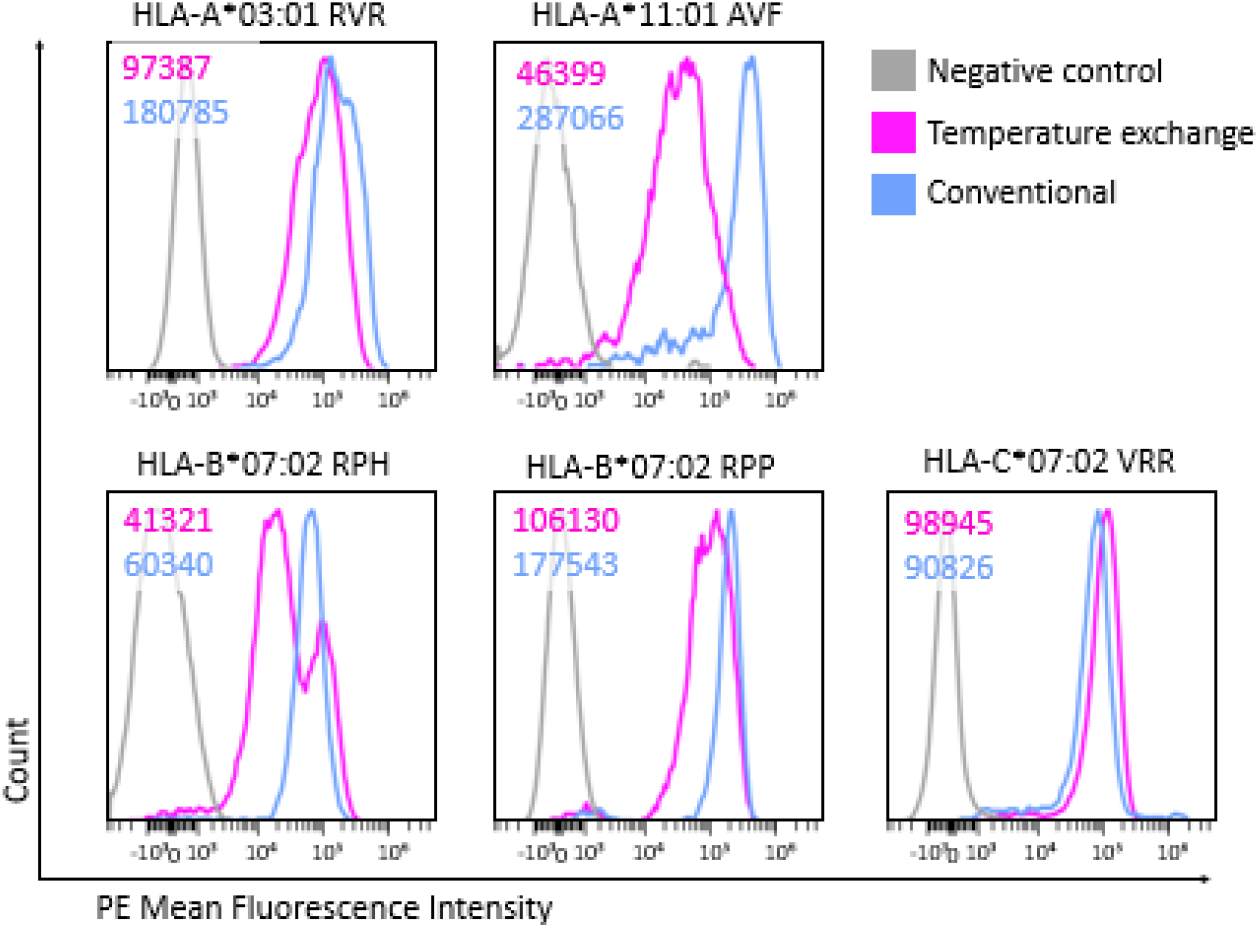
Comparison of temperature exchange multimers to conventional multimers on clonal T cell lines. Clonal CD8+ T cell lines stained with 13.3 ng/μl of PE-conjugated temperature exchange tetramers (pink) or with conventional tetramers (blue). The specificities of the T cell lines are depicted with the HLA-type and first three amino acids of the peptide for each line. Y-axis shows the individually scaled counts and x-axis shows mean fluorescent intensity of PE.

### Detection of virus-specific CD8+ T cells in peripheral blood

To evaluate the practical utility of our additional HLA alleles for temperature-based peptide exchange MHC-I multimers using human peripheral blood samples, we compared the performance of temperature-based peptide exchanged multimers to that of classical multimers within an immune monitoring framework. This is particularly relevant in the context of monitoring virus-specific T cells in immunocompromised patients, such as those undergoing T cell-depleted allogeneic stem cell transplantation and adoptive cell therapy. Herpesviruses, including human cytomegalovirus (HCMV) -and Epstein-Barr virus (EBV), can cause severe infections in these patients. Restoring T cells is essential to prevent these infections and to minimize complications and mortality. We analyzed peptides derived from HCMV and EBV for the HLA types HLA-A*11:01, HLA-B*07:02, and HLA-C*07:02 in peripheral blood samples from healthy donors. Additionally, we included the previously developed temperature-based peptide exchange multimers for HLA-A*02:01 in this analysis[18]. We exchanged PE- and APC-labeled multimers for a selection of these peptides to measure T cell frequencies in peripheral blood mononuclear cells (PBMCs) and compared them to traditional folded multimers. The antigen-specific T cells identified with temperature-exchanged multimers stained a distinct cell population (Figure 5). Importantly, we did not observe any MHC-I multimer positive T cells in two irrelevant donors (Supplemental Figure 5). The percentages observed are comparable to those observed with conventional multimers (Figure 5). The peptides are listed in Supplemental Table 2 and the gating strategy of the immune monitoring framework is shown in Supplemental Figure 6. These result illustrates the efficiency and versatility of our technology in rapidly generating diverse MHC-I multimers from a ready-to-use stock, enabling the detection of antigen-specific T cells, even at the low frequencies commonly observed in primary immune monitoring samples.

**Figure 5:**
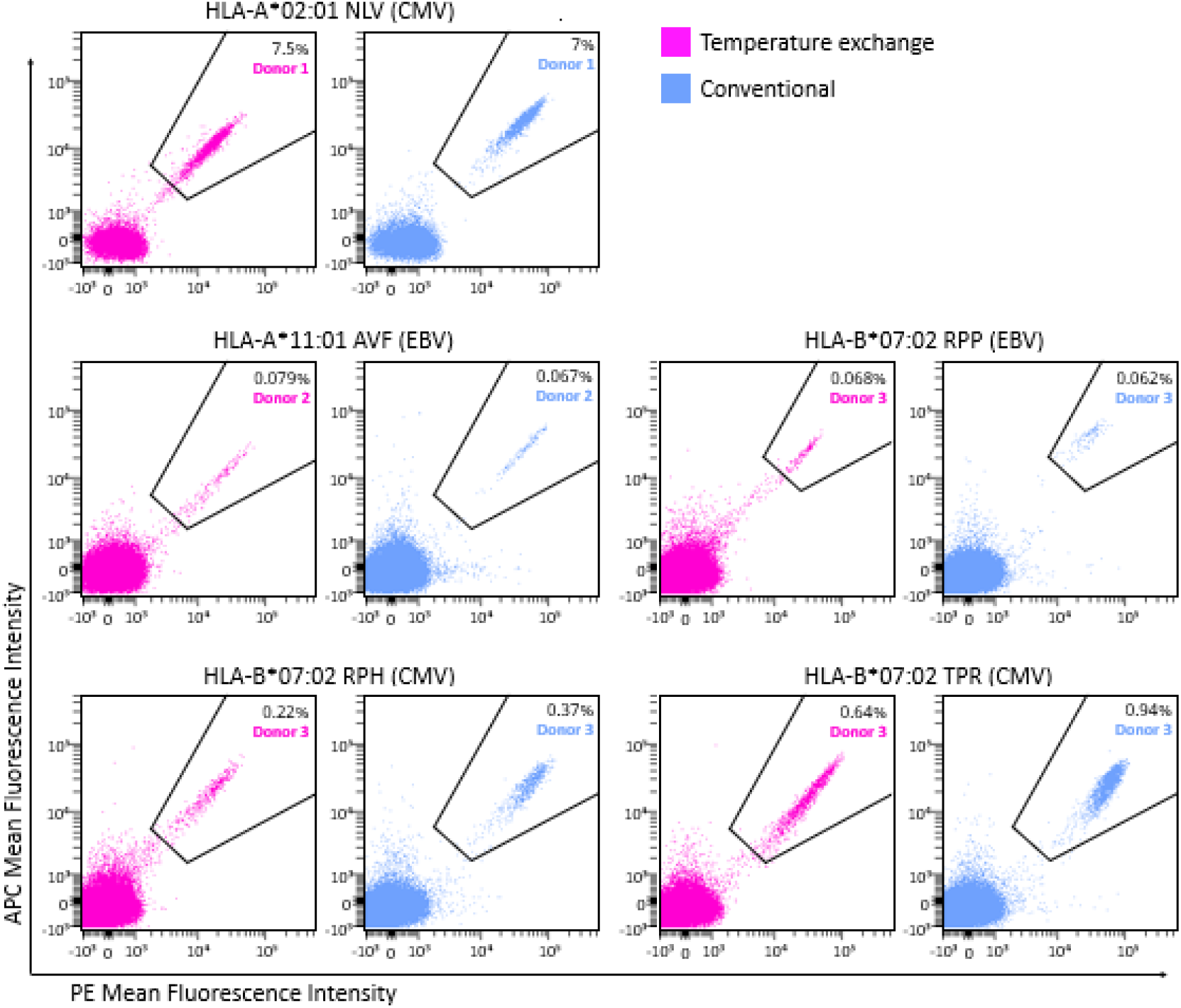
Comparison of temperature exchange tetramers to conventional tetramers on PBMCs. PBMCs of healthy individuals were incubated with peptide exchange (pink) or conventional (blue) tetramers. Donor PBMCs were CMV- or EBV-positive based on serology and were incubated with the relevant tetramer based on donor HLA-type. Tetramers were conjugated to APC (y-axis) as well as PE (x-axis). Gate shows the tetramer-binding CD8+ T cells and the percentage of total CD8+ T cells is depicted in the top right corner.

## Discussion

The use of multimeric versions of peptide MHC-I complexes has become an important technology to detect and quantify antigen-specific CD8+ T cells[29-32]. To identify antigen-specific T cells using flow cytometry, fluorescently labeled MHC multimers are widely employed. There is an increasing interest in the detection of CD8+ T cells immune responses since they are important for the protection against pathogens or cancer and can be used to define antigens for vaccination purposes.

Two challenges in advancing this field are the need for high-throughput approaches and the lack of ready-to-use reagents. Classical peptide-MHC-I complexes are each produced in a separate production run, required for each unique peptide-HLA complex. The process is elaborate, time consuming and demands highly skilled people. This restricts the generation of a larger library of peptide-HLA combinations and thereby results in incomplete detection of the full CD8+ T cell response against an antigen. Previously, it has been shown that peptide-exchange technologies, such as UV-based peptide exchange, are able to upscale towards higher throughput approaches. While the stability of UV-exchanged pMHC complexes can be assessed using methods such as thermal shift assays and size-exclusion chromatography, the efficiency of peptide exchange is more difficult to quantify directly. Temperature-based peptide exchange is controlled, simple, fast, and quantitative if the correct conditions are identified, as for the HLA alleles in this manuscript. Moreover, the broader availability and simplicity of equipment needed for thermal exchange compared to UV sources enables wider adoption of this approach. The time to prepare the materials for flow analyses is around 1 hour and consists of taking the exchange MHC-I multimer from the fridge and warming it up in the presence of peptides of interest. This can be immediately followed by staining the CD8^+^ T cells for flow cytometric analysis.

Additionally, MHC-I molecules can interact with other surface proteins in both a peptide-dependent and peptide-independent manner. For example, they bind to the inhibitory receptor Leukocyte immunoglobulin-like receptor subfamily B member 1 (LILRB1), independently of the sequence of the presented peptide, while interactions with receptors such as Killer Immunoglobulin-like Receptors and CD55 are peptide-dependent[33-36]. Consequently, a positive flow cytometry signal does not necessarily indicate interaction with the TCR of CD8+ T cells. It is therefore essential to exercise caution when interpreting MHC-I multimer results to ensure accurate conclusions. These diverse MHC-I based interactions may provide further utility for studying various immune cell subsets beyond T cells, including natural killer (NK) cells and other innate immune cells from a peptide MHC-I specific perspective.

We have set out to determine whether temperature-based peptide-exchange technology can be developed into a broadly applicable platform for the screening of human CD8+ T cells. For this purpose, we designed conditional peptide ligands for four different human MHC-I alleles across the three human HLA loci: HLA-A*03:01, HLA-A*11:01, HLA-B*07:02 and HLA-C*07:02. Refolding reactions with the temperature-based peptide are efficient and result in stable peptide MHC-I complexes at 4°C, which are unstable with increased temperature. The presence of an high-affinity peptide during temperature-based peptide exchange ensures the replacement of the temperature conditional ligand in the MHC-I peptide-binding groove. This leads to the generation of the peptide-MHC-I complex with a desired antigen specificity. Importantly, we show that the temperature-based peptide exchange can be performed using the multimerized and fluorescently labeled MHC-I complexes. This allows the development of ready-to-use MHC-I multimers, with high-throughput abilities demonstrating the ease of use of temperature-based peptide exchange technology.

The temperature-based peptide exchange method for generating MHC-I multimers, has several limitations. First, effective peptide exchange necessitates the design of HLA-specific ligands, adding complexity compared to approaches that utilize empty-loadable MHC-I multimers. This reliance on specific ligands could also affect the specificity and functionality of T cell responses, particularly when low-affinity peptides are involved. Additionally, although the method allows for rapid ready-to-use multimer generation, the inherent stability of some MHC-I complexes could limit the types of peptides that can be successfully exchanged. Furthermore, while this system is designed to be accessible to a wide range of laboratories, the initial need for appropriate peptide selection and optimization might still pose a challenge for some users. This technology is most likely not optimal for low affinity peptides, as the template peptide itself must also have low affinity. Lastly, despite the benefits of temperature-based peptide exchange technology, there is still a risk that some combinations may not provide the desired specificity and functionality. This highlights the need for ongoing refinement and validation of the method across different MHC-I alleles. Additionally, using complementary peptide exchange methods may enhance overall effectiveness as has been done for UV-exchange multimers and empty disulfide-stabilized multimers[37].

Current peptide exchange technologies predominantly target HLA alleles common in individuals of Western European descent. However, we have started expanding this work to include HLA-A*11:01, which is most prevalent in East and Southeast Asian populations [38, 39]. Currently there are temperature-based peptide-exchange multimers for six HLA alleles, namely HLA-A*02:01, HLA-A*03:01, HLA-A*11:01, HLA-B*07:02, HLA-C*07:02 and HLA-E[19, 20]. With this set of MHC-I reagents, a deep analysis could be possible for people with an European genetic background.

A variety of MHC-I reagents are currently available for use in CD8+ T cell diagnostics, including classical folded MHC-I multimers, empty stabilized multimers[9] and UV-exchange multimers. These MHC-I reagents are being used to determine virus specific T cell diagnostics, personalized neoantigen-specific T cell diagnostics and more. We propose that the ready-to-use temperature-based peptide exchange MHC-I technology could further advance the field, as these reagents simplify the process by eliminating additional steps typically required for MHC-I multimers production.

## Material and methods

### PBMCs isolation

PBMCs were isolated from healthy donors and obtained via Sanquin or biobanked by the Department of Hematology (LUMC) after informed consent, in accordance with the Declaration of Helsinki. PBMCs were isolated from buffy coats or whole blood using Ficoll-Isopaque and used directly or stored using cryo-preservation. Biobanked PBMCs were thawed in Iscove Modified Dulbecco Medium (IMDM; Lonza, Basel, Switzerland) supplemented with 10% heat-inactivated fetal bovine serum (FBS; Sigma-Aldrich, Saint Louis, MO, USA), 2.7 mM L-glutamine (Lonza), 100 U/mL penicillin (Lonza), and 100 μg/mL streptomycin (Lonza; 1% p/s), and subsequently treated with 1.33 mg/mL DNAse to minimize cell clumping. PBMCs were either used for isolation of clonal T cell lines or for flow cytometry analysis.

### Clonal T cell lines

Clonal T cell lines were isolated and expanded as previously described[40, 41]. In brief, PBMCs were incubated with peptide-HLA multimers and antibodies targeting CD14, CD8 and CD4 to isolate CD14-CD8+CD4-multimer-binding cells using fluorescence-activated cell sorting (FACS). In some cases, cells were pre-selected prior to FACS using magnetic-activated cell sorting (MACS) based on peptide-HLA multimer-binding. Obtained cells were further expanded either as bulk[40] or single cell per well[41], in the presence of irradiated feeders with or without irradiated Epstein-Barr virus lymphoblastoid cell lines (EBV-LCLs), phytohemagglutinin (PHA) or specific peptides. See Supplemental Table 1 for T cell line details.

### Flow cytometry

1-2×10^6^ PBMCs or 0.05-0.1×10^6^ T cells were seeded in a round bottom 96-wells plate. Cells were stained with the viability dye Zombie-Red (Biolegend, San Diego, CA, USA) for 25 min at room temperature after which the cells were washed in PBS containing 0.8 mg/mL albumin (FACS buffer). Cells were stained with 20 μl multimer staining mix, incubated for 30 minutes at 4°C. Conventional multimers were generated in-house, as previously described[40]. After washing, cells were stained with 20 μl of an antibody cocktail containing: αCD8-pacific blue (cat#558207, BD Biosciences), αCD4-BV510 (cat#562970, BD Biosciences), αCD14-FITC (cat#555397, BD Biosciences), αCD19-FITC (cat#555412, BD Biosciences) and αCD16-FITC (cat#335035, BD Biosciences). After incubation for 20 minutes at 4°C, cells were washed and measured on a 3-laser Aurora (Cytek Biosciences) or LSR II (BD Biosciences) flow cytometer. Flow cytometry data analysis was performed using OMIQ.ai. See Supplemental Table 2 for patient details regarding CMV- and EBV status and HLA typing.

K562 LILRB1^OE^ cells were harvested, washed and 100K cells were stained in 50 uL facs buffer (1% FCS in PBS) supplemented with multimers (1:80 dilution for temp exchange or 200 ng conventionel tetramer) or anti-LILRB1 FITC (1:40, clone REA998, miltenyi biotec) for 30 minutes on ice shielded from light. After 2 rounds of washing, cells were measured in 100 uL facs buffer on a BD FACS LSR II 4L Full (BD Biosciences, San Jose, CA, USA) flow cytometer and data was analyzed using FlowJo FlowJo (V10.8.1).

### Peptide synthesis and purification

Peptides were synthesized at the LUMC Peptide Facility by standard solid-phase peptide synthesis using Syro I and Syro II synthesizers. Peptides were purified using reversed-phase HPLC (Waters) using a mobile phase of water/acetonitrile gradient containing 0,1% TFA on a C18 column (X-bridge, Waters). Purity of the peptides was confirmed using LC-MS (Micromass LCT Premier; Waters).

### Generation of temperature sensitive monomers and multimerization

Temperature-based sensitive monomers were produced as previously described[21]. Briefly, recombinant HLA-A/B/C heavy chain and B2M light chain were produced as inclusion bodies in bacterial strain BL21 (DE3). These purified inclusion bodies containing 2.5 mg of heavy chain were solubilized in 8 M urea and folded for 5 days at 10°C with 1.2 mg prefolded B2M and 3 mg temperature-sensitive peptide (A3: RGAVRATKR; A11: RFKMFPEVR; B07: RKELVRPAL; C07: IKYNYPAMA) in 50 ml folding buffer (400 mM L-Arginine, 0,5 mM oxidized glutathione, 5 mM reduced glutathione, 2 mM EDTA, 100 mM Tris.HCl pH 8, glycerol 5% and half a tablet of protease inhibitor cocktail (complete, Roche)). Temperature-based sensitive monomers were concentrated on a 30 kDa filter (Amicon Ultra-15) and biotinylated with BirA ligase overnight at 4°C. Biotinylated Temperature-based sensitive monomers were purified with size exclusion chromatography (Superdex 75 10/300 GL) on a NGC system (Bio-Rad). Properly folded complexes were concentrated on a 30 kDa filter (Amicon Ultra-15) and the concentration was measured on a NanoDrop (Thermo Fisher Scientific) and stored in 300 mM NaCl and 20 mM Tris-HCl, pH 8.0 with 15% glycerol at -80°C. Next, the monomers were multimerized with PE-labeled streptavidin and/or APC-Labeled streptavidin (SA-PE, SA-APC; Thermo Fisher Scientific, Invitrogen) to a final concentration of 0,625 μM. For peptide exchange, temperature-based monomers were incubated at a final concentration of 0,5 μM and 50 μM of selected peptide. This was diluted 1:20 with PBS to be added to CD8+ T cells.

### HPLC analysis of temperature-mediated peptide exchange

To initiate temperature-based peptide exchange, a 0.5 μM peptide–MHC-I complex was mixed with 50 μM exchange peptide in 110 μl PBS and incubated under defined conditions. After centrifugation at 14,000 g for 1 min, the supernatant was analyzed using size-exclusion chromatography on a Shimadzu Prominence HPLC system with a 300 × 7.8 mm BioSep SEC–s3000 column and PBS as mobile phase. Data were analysed using Shimadzu LabSolutions software.

### LILRB1 over-expressing K562 cell line

The ECD of LILRB1 was fused to the platelet-derived growth factor receptor transmembrane spanning domain and cloned into pCDH (ires puro) vector (deposited at Addgene). Virus production, HEK 293T cells were transfected with packaging plasmids pMDLg/pRRE (Addgene), pRSV-Rev (Addgene), pCMV-VSV-G (Addgene) together with the d_LILRB1 plasmid using polyethyleneimine (Polyscience Inc.) . Virus was harvested, filtered and K562 cells were transduced with LILRB1 in the presence of 8 μg/ml polybrene (Millipore). Transduced cells were selected using puromycin (2 μg/ml, Thermo Fisher) and sorted twice based on high LILRB1 antibody staining (clone REA998, Miltenyi Biotec, Fitc) on a CytoFLEX SRT Benchtop cell sorter (Beckman Coulter, Brea, CA, USA). K562 cells transduced with LILRB1 were cultured in Iscove’s Modified Dulbecco’s Medium (IMDM) supplemented with 10% FBS (Bodinco), 1.5% glutamine (200⍰mM; Lonza) and 1% penicillin/streptomycin (200⍰mM; Lonza). The stability of PE fluorescent tetramers was indicated by PE-positive K562 cells[27, 36].

## Glossary

MHC Class I; Peptide Exchange; HLA Alleles; CD8+ T Cells; Temperature-sensitive Multimers; Antigen-specific Immunity; Fluorescent Labeling; High-throughput Technology

## Acknowledgements

We thank Dris el Atmioui and Cami Talavera Ormeño (Department of Chemical Immunology, LUMC, Leiden, Netherlands) for synthesis of peptides. We would also like to thank the flow core facilities of the LUMC, Leiden, The Netherlands. This work was supported by a LUMC strategic grant and starters grant from NWO to F.A. Scheeren.

## Supplemental Figure legends

**Supplemental figure 1: Thermal stability of template peptide with MHC-I monomers**. Thermal stability of the template peptide MHC-I complex at 50°C using analytical size exclusion HPLC for HLA-A*03:01, HLA-A*11:01, HLA-B*07:02 and HLA-C*07:02 (chromatograms). Input, no peptides and plus high-affinity peptide. One of at least three representative experiments is shown.

**Supplemental Figure 2: Thermal stability optimization of the template peptide MHC-I monomers**. Thermal stability of template peptide MHC-I complex to determine optimal temperature with one high-affinity peptide for all alleles. Primary data of temperature based peptide exchange analyzed by gel filtration chromatography at indicated temperatures. Timing is 1 hours, unless other indicated. The exchange peptides were used are depicted in Table 1. One of at least three representative experiments is shown.

**Supplementary figure 3: Dilution of temperature exchange multimers on T cell lines**. Clonal CD8+ T cell lines stained with PE-conjugated temperature exchange multimers that either contained an irrelevant (negative control; grey) or a relevant peptide (pink) in different concentrations. The different concentrations are depicted in three different shades of pink (1:6, :1:8 and 1:4). The specificity of the T cell clones are depicted above each figure as the HLA-restriction and first three amino acids of the peptide. Y-axis shows the individually scaled counts and x-axis shows mean fluorescent intensity of PE.

**Supplementary figure 4: Comparison of temperature exchange (TE) multimers to conventional multimers at different concentrations**. Clonal CD8+ T cell lines were unstained (negative control; grey) or stained with PE-conjugated temperature exchange multimers (pink) or with conventional multimers (blue). The specificity of the T cell lines are depicted on the left side as the HLA-restriction and first three amino acids of the peptide. The concentration of the multimers is depicted above. Y-axis shows the individually scaled counts and x-axis shows mean fluorescent intensity of PE.

**Supplementary figure 5: Comparison of temperature exchange multimers to conventional multimers on irrelevant PBMCs**. PBMCs of healthy individuals were incubated with peptide exchange (pink) or conventional (blue) multimers. The peptide-HLA combinations of the multimers and from which virus the peptide is derived, are depicted above each figure. Each figure is an overlay of two donors that were CMV- or EBV-positive based on serology and had an irrelevant HLA-type for the multimers. Multimers were conjugated to APC (y-axis) as well as PE (x-axis). Gate shows the multimer-binding CD8+ T cells and the frequency of total CD8+ T cells is depicted in the top right corner.

**Supplementary figure 6: Flow cytometry gating strategy example**. Representative example of gating strategy used for flow cytometry. A) Gating strategy used for CD8+ T cell clones. All events were gated on lymphocytes and single cells. B) Gating strategy used for PBMCs. All events were gated on lymphocytes, single cells, live cells, CD14-CD19-CD16-, CD8+CD4- and multimer+.

**Supplemental Table 1: Clonal** CD8+ T cell line details

**Supplemental Table 2:** Donors used for detection of virus-specific CD8+ T cells using temperature-based sensitive multimers

**Supplemental Table 1:**
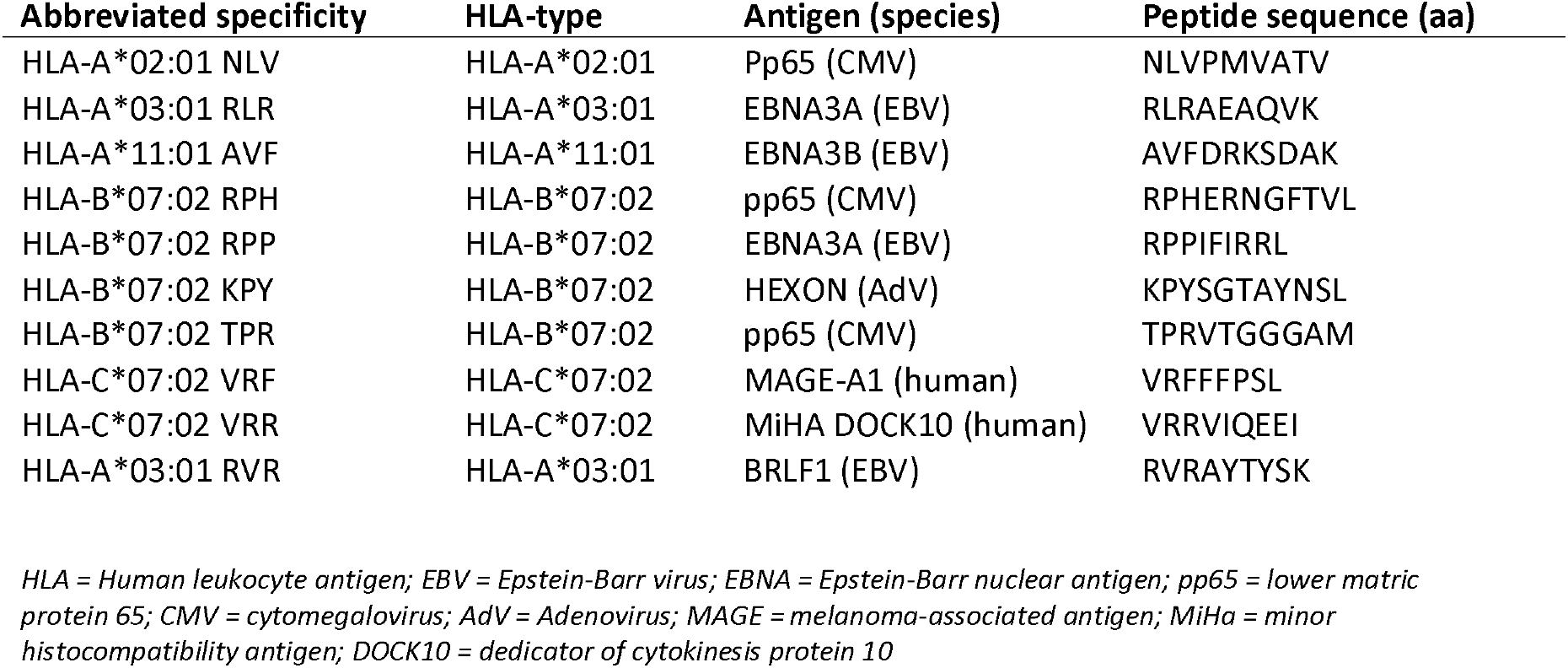
**Clonal** CD8+ T cell line and exchange peptide details

**Supplemental Table 2:** Donors used for detection of virus-specific CD8+ T cells using temperature-based sensitive multimers

| Donor # | CMV-status | EBV-status | HLA-A* | HLA-B* | HLA-C* |
| --- | --- | --- | --- | --- | --- |
| 1 | + | + | 02:01 | 44:02 | 05:01 |
| 2 | + | + | 01:01 | 08:01 | 03:01 |
|  |  |  | 11:01 | 15:01 | 07:01 |
| 3 | + | + | 01:01 | 07:02 | 07:01 |
|  |  |  | 03:01 | 08:01 | 07:02 |

