## supplemental figures for "Temperature-based MHC class-I multimer peptide exchange for human HLA-A, B and C"

Supplemental Figure 1

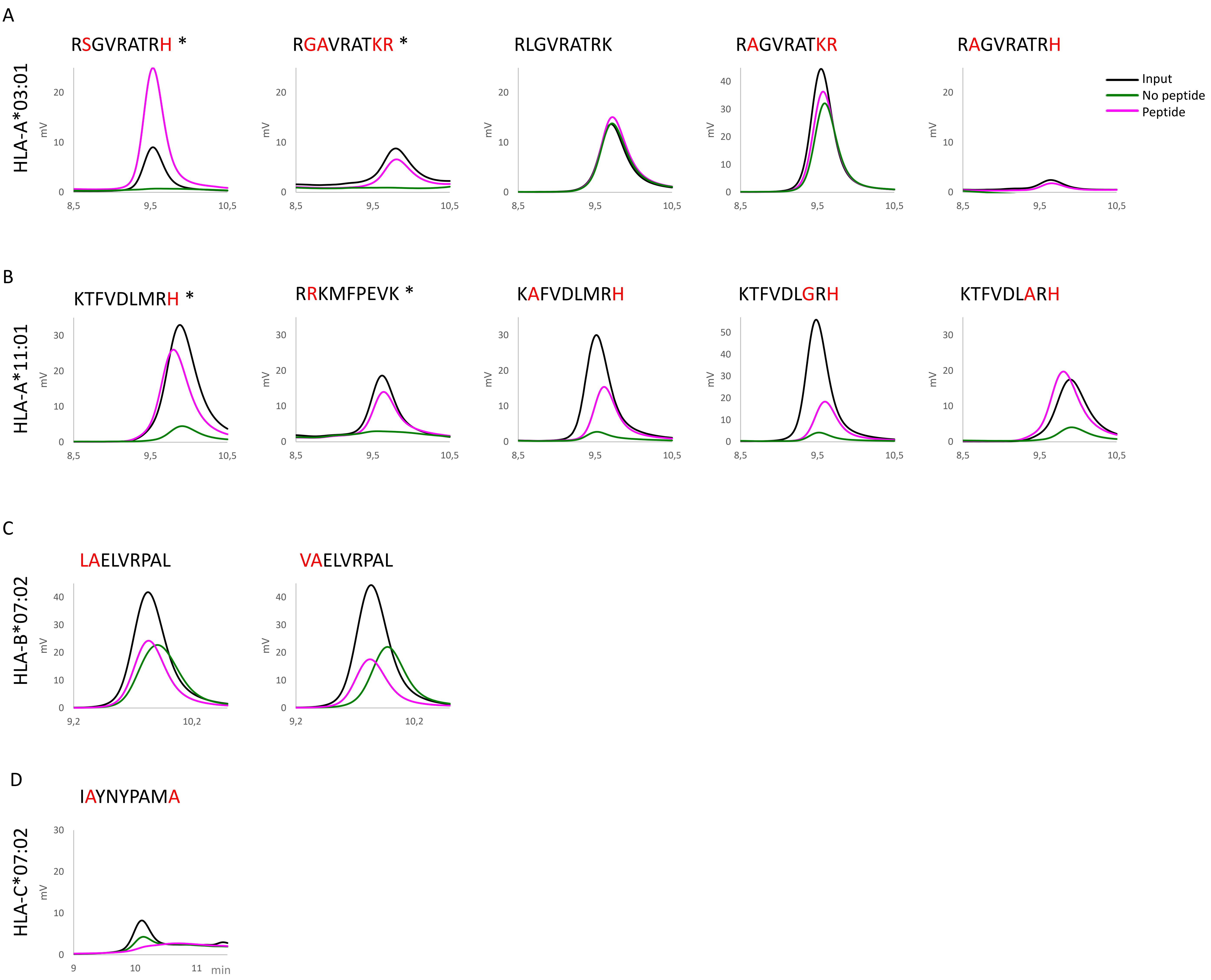

Supplemental Figure 2

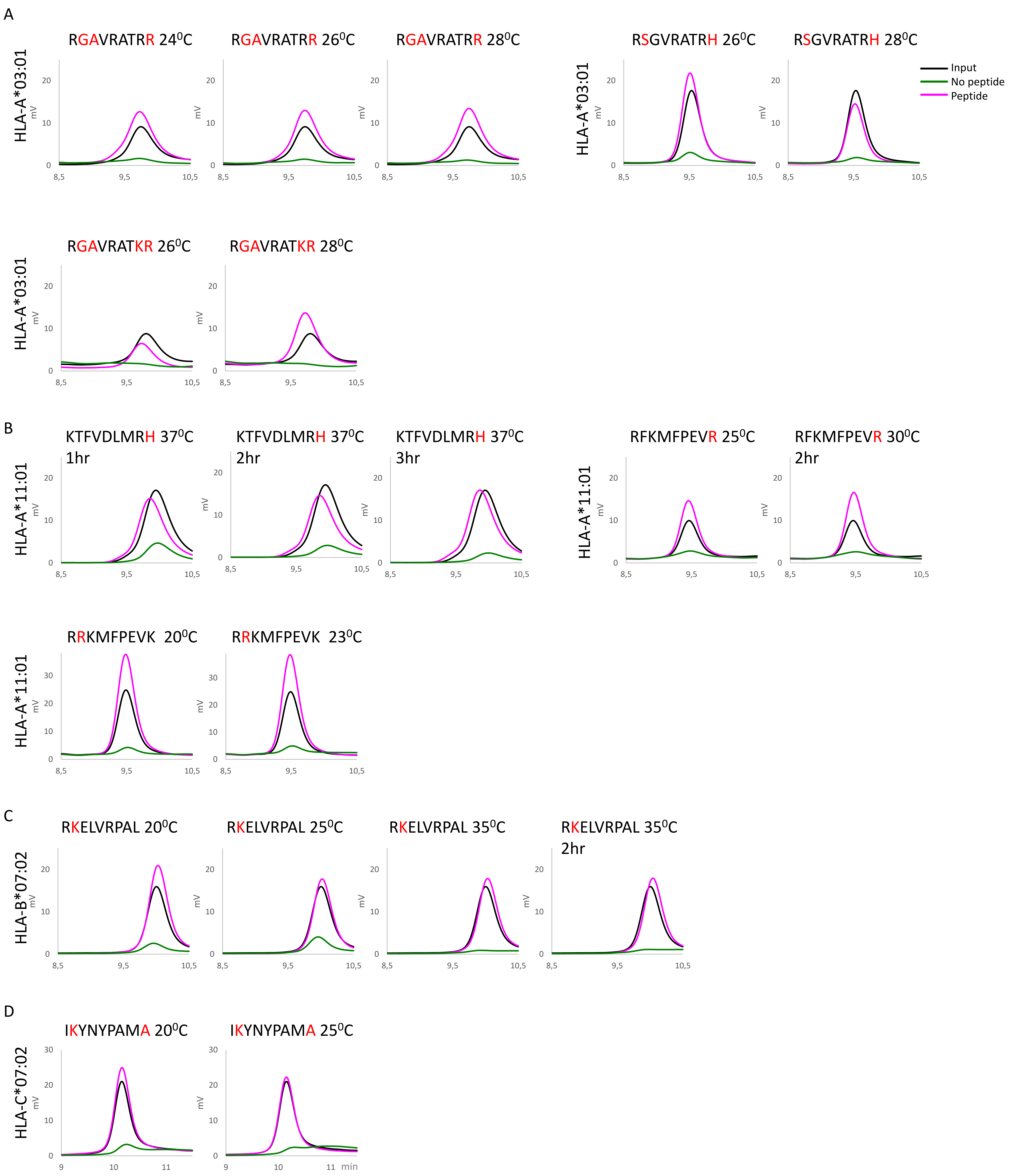

Supplemental Figure 3

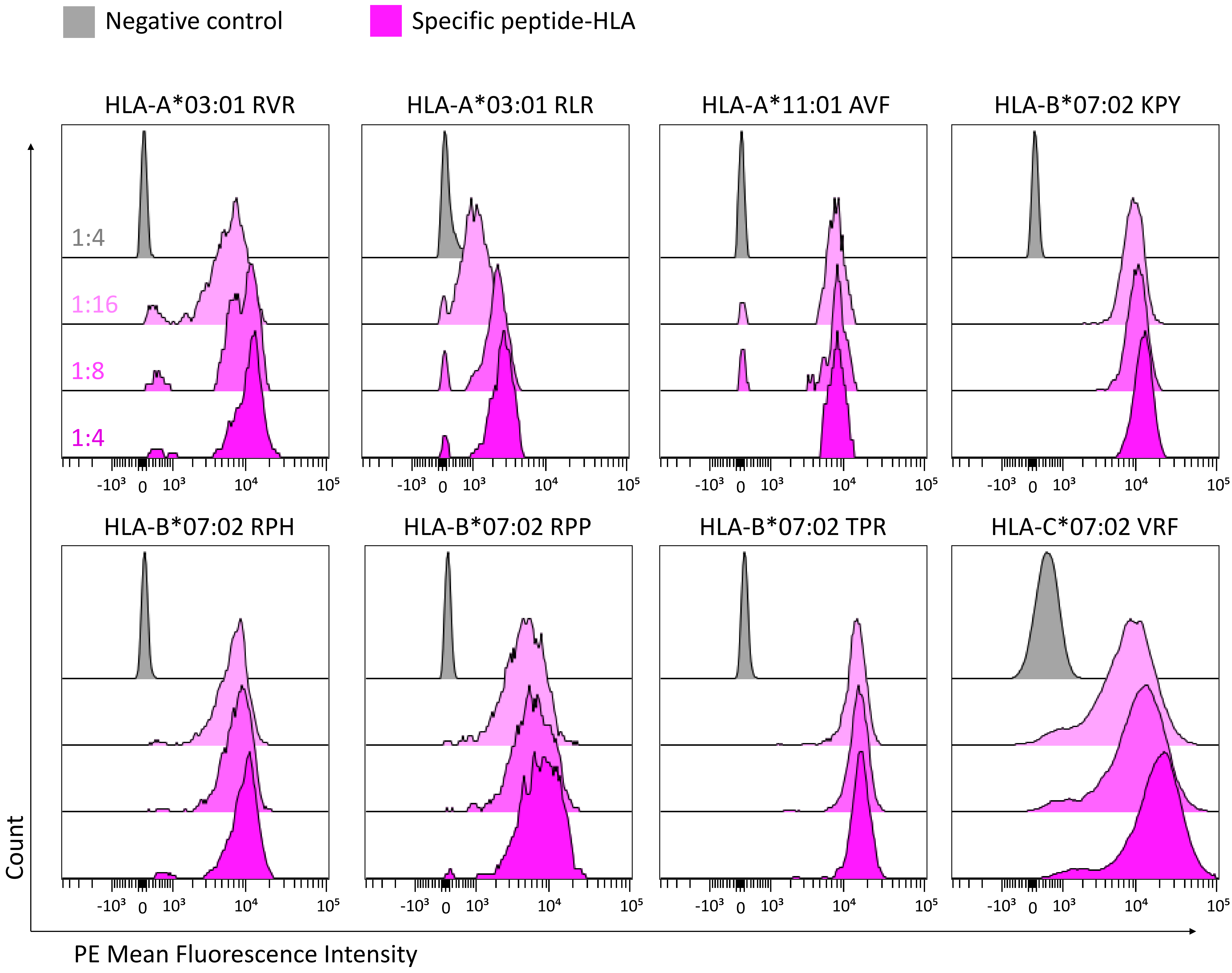

Supplemental Figure 4

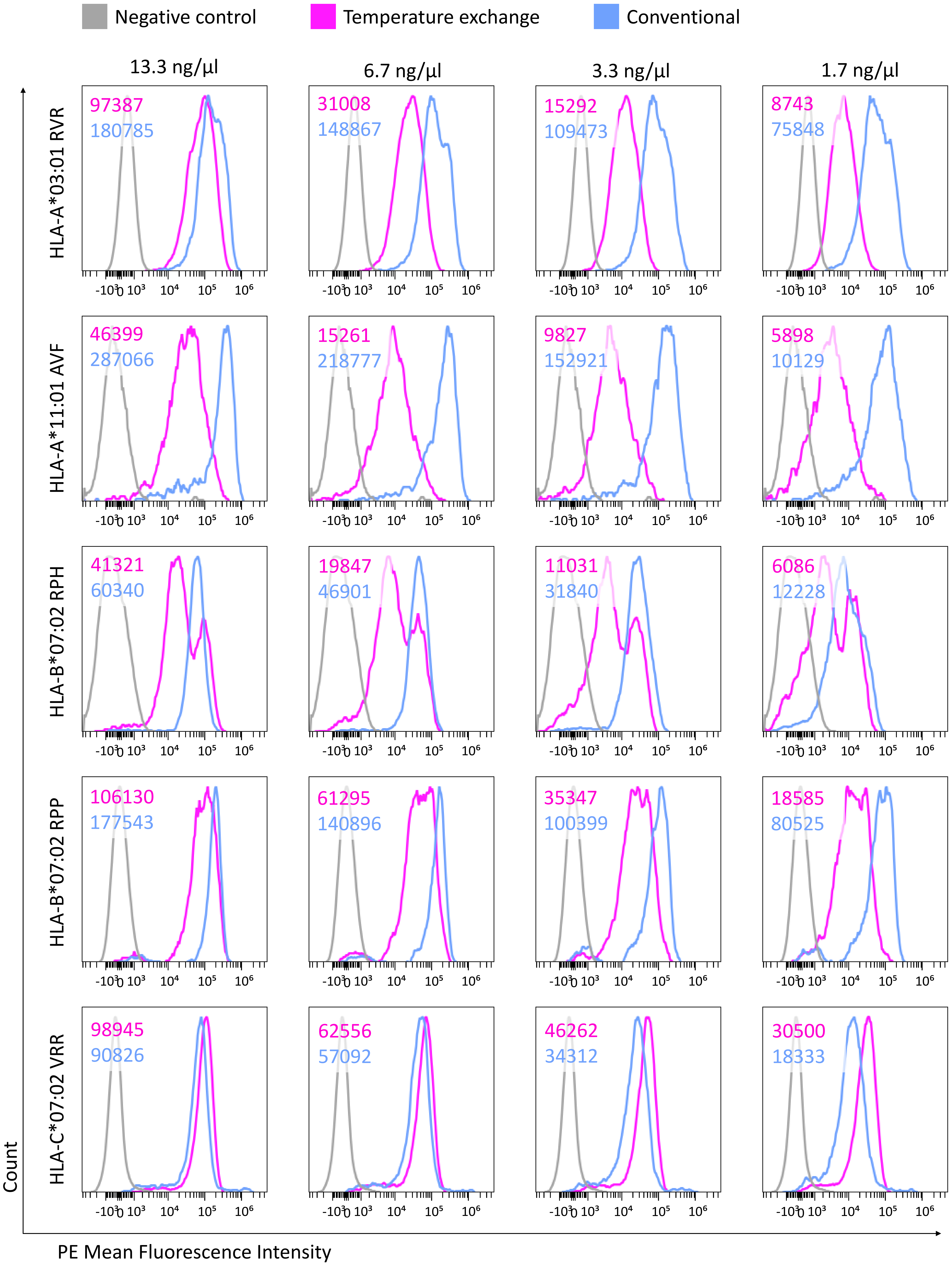

Supplementary figure 5

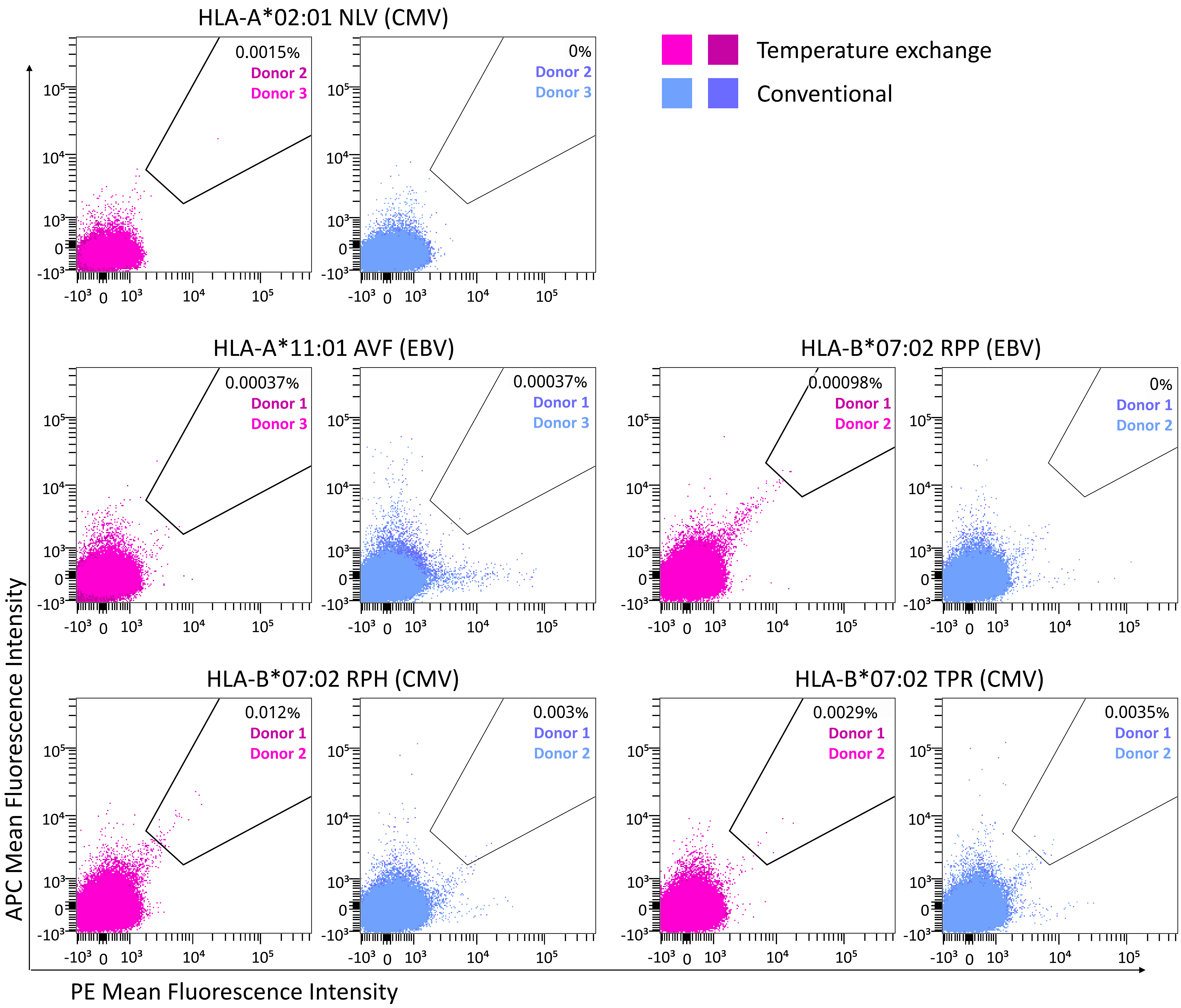

Supplementary figure 6

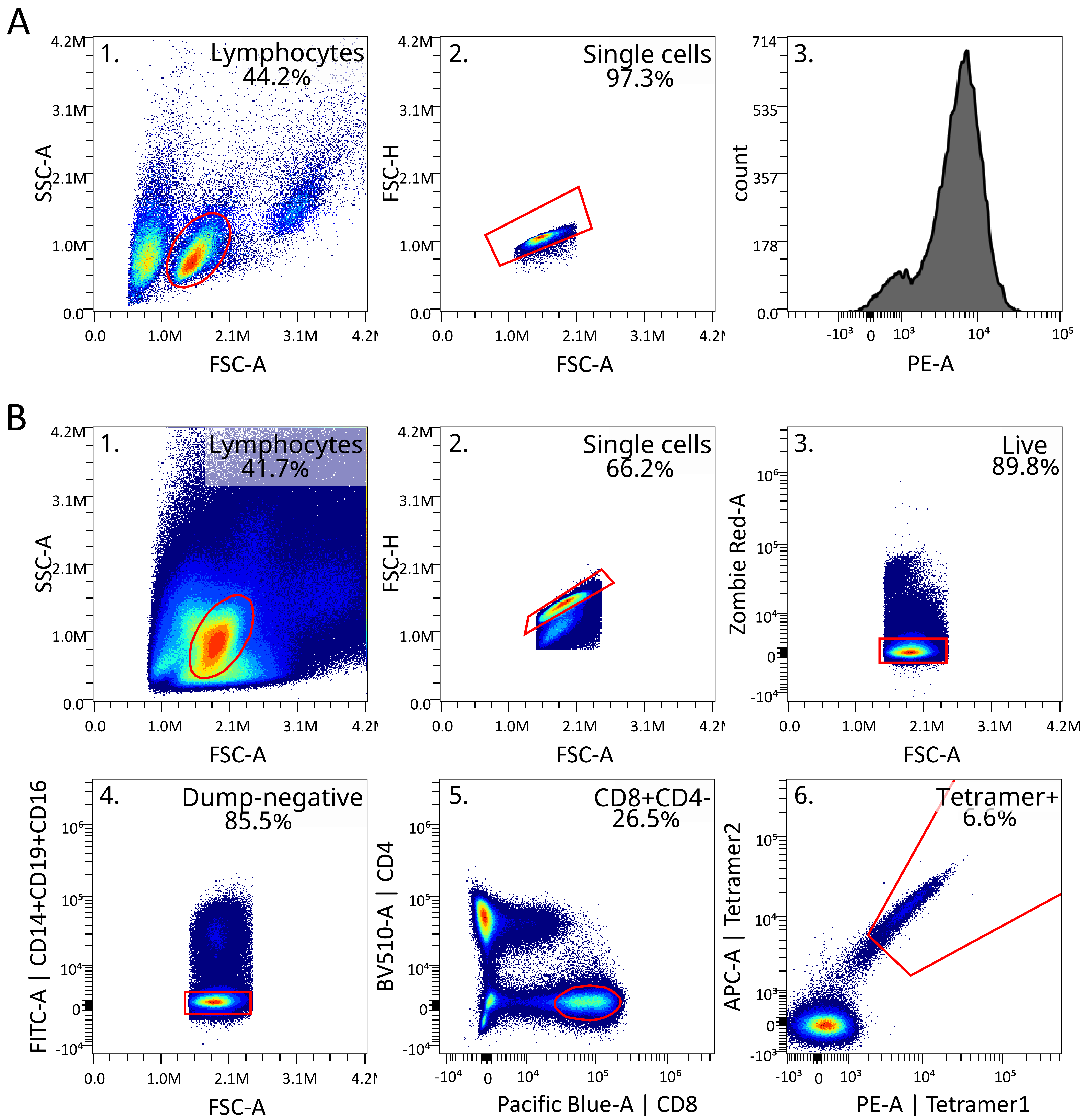
